# A Pipeline for Predicting Variant Associations with Complex Disorders via Functional Annotations

**DOI:** 10.64898/2026.10.21.760618

**Authors:** Francesco Gualdi, Zeno Darani, Daniele Malpetti, Marco Scutari, Francesca Mangili

## Abstract

The study of complex disorders is limited by the lack of structured data sets that integrate genetic association signals with rich functional annotations. While genome-wide association studies (GWAS) provide large collections of associated variants, these are heterogeneous, study-dependent, and not directly suitable for machine learning applications. Although frameworks such as Combined Annotation Dependent Depletion (CADD) offer comprehensive variant-level features, there is no standardised procedure to construct curated, balanced, and reproducible data sets for the systematic study of complex traits.

We introduce a scalable and reproducible pipeline for generating CADD-annotated variant data sets associated with complex disorders defined via Experimental Factor Ontology (EFO) terms. The pipeline integrates GWAS summary statistics, genome mapping, chromosome-stratified negative sampling, and large-scale annotation, while addressing class imbalance and feature redundancy. The resulting data sets provide curated, balanced, and annotation-rich representations of variant–trait associations, suitable for statistical learning, systematic model evaluation and comparison, and analysis of functional annotation contributions and interactions.

## 1 Introduction

Deciphering the genetic architecture of complex disorders remains a central challenge in human genomics. Unlike Mendelian diseases, which are often driven by single high-impact variants, complex traits arise from the combined effects of many variants, each typically exerting a modest influence [1].

Large-scale efforts such as Genome-Wide Association Studies (GWAS) [2] have successfully identified thousands of loci associated with complex traits. However, translating these associations into biological insight remains nontrivial. Most associated variants lie in non-coding regions, and only a small fraction have a clear mechanistic interpretation. As a result, there is a growing need for computational frameworks that can prioritise and annotate variants based on their likely functional impact [3].

One widely adopted approach is the use of integrative scoring systems such as Combined Annotation Dependent Depletion (CADD) [4], which aggregates diverse genomic annotations, including conservation metrics, regulatory features, and functional predictions, into a single score reflecting a variant’s deleteriousness.

Extensive validation of variant association prediction frameworks has been performed for rare Mendelian diseases, particularly using variants curated in ClinVar. These efforts have demonstrated the utility of computational predictors in distinguishing associated or non-associated variants in a monogenic context, with more recent advances implementing large-scale foundational models to further improve predictive performance [5].

In contrast, comparatively little work has focused on evaluating such approaches for variants associated with complex disorders, as collected in the GWAS Catalog [2]. Existing studies have made only moderate progress in assessing variant association in this setting, often without anchoring predictions to specific diseases [6]. Moreover, assessing variants in isolation is inherently limited for complex traits, where genetic risk is distributed across many loci with small individual effects.

In this work, we present a scalable and reproducible pipeline for generating curated data sets of CADD annotation-enriched variants associated with a user-defined selection of complex disorders, as defined by EFOs [7]. The pipeline integrates publicly available association data with automated annotation workflows to generate structured data sets suitable for downstream statistical analyses and for benchmarking machine learning methods. By systematically combining genetic association evidence with functional annotations, this pipeline provides a customizable reference data set for evaluating variant effect prediction tools in the context of complex disorders, ultimately contributing to a better understanding of the regulatory genomic mechanisms underlying these diseases.

We demonstrate the presented pipeline through a use case on predicting variants associated with Coronary Artery Disease (CAD) and compare the performance against that of CADD deleteriousness scores.

## 2 Pipeline

Figure 1 summarizes the steps of the pipeline, which takes as input a collection of disorders represented by EFO codes together with their associated GWAS studies identified by GWAS Catalog Study (GCST) IDs, and produces a balanced, annotated variant dataset linked to the condition of interest, ready for downstream machine learning applications.

**Figure 1.**
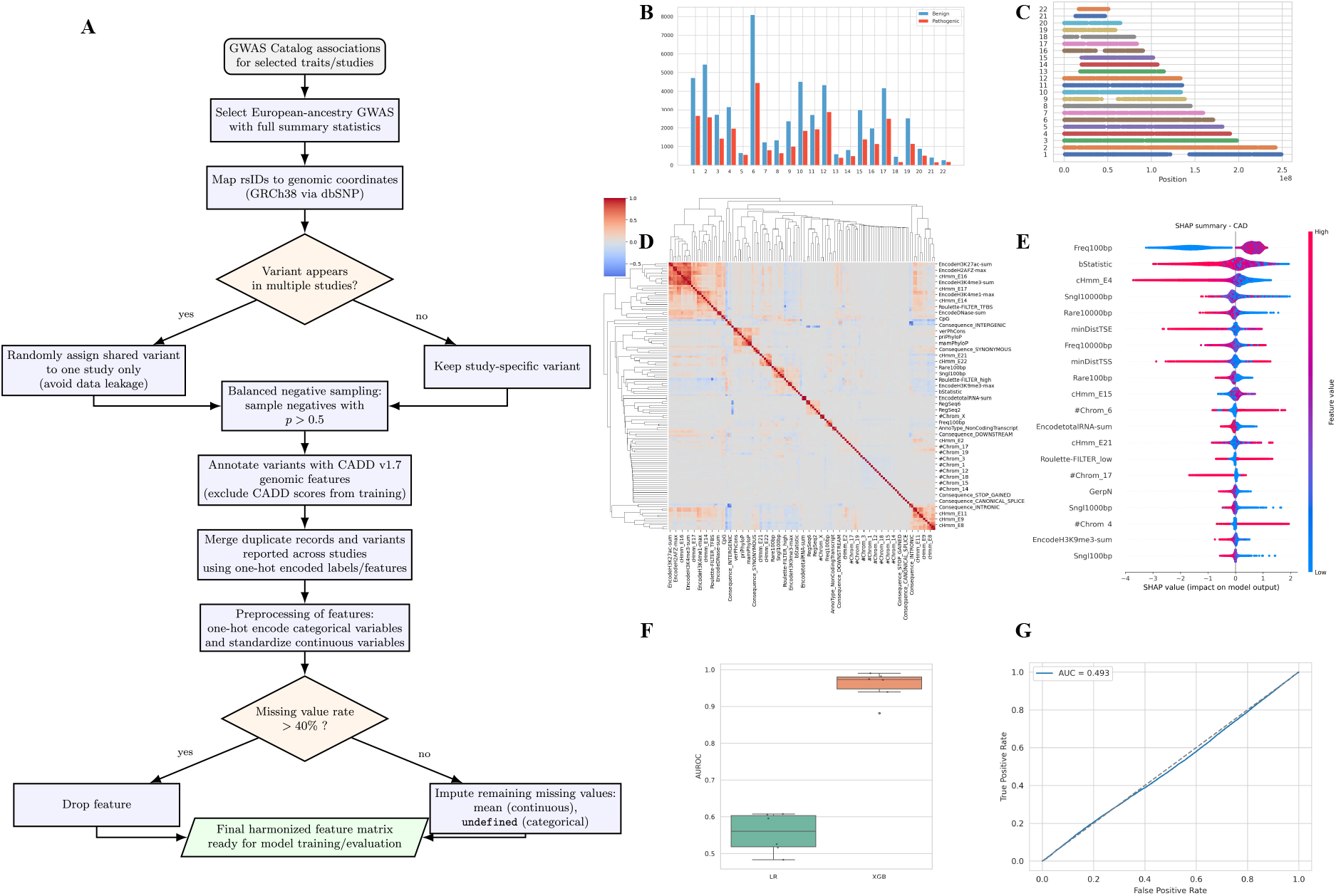
Data processing pipeline and performance evaluation for CAD GWAS variant classification. (A) Flowchart of the data curation workflow, fromGWAS selection to the generation of the data set. (B–C) Distribution of variants across chromosomes 1–22, shown by frequency and genomic position. (D) Hierarchical clustering heatmap showing the correlation structure between CADD-annotated genomic features. (E) SHAP summary plot illustrating the global impact of top features on model predictions on CAD, where colour represents feature value. (F–G) XGBoost and Baseline LR mean AUC across CAD studies, AUROC Computed using Raw Scores from CADD (AUC ≈ 0.5).

### GWAS

The input consists of a Python dictionary mapping EFO trait codes to their associated GCST IDs. An optional parameter allows filtering by ancestry. The pipeline queries the GWAS Catalog for studies with full summary statistics available and stores the data locally.

### rsID mapping

After generating a list of the variants contained in the data, we map their rsIDs to the GRCh38 reference genome using dbSNP [8]. To prevent data leakage and ensure unbiased performance estimates at the study level, when a variant was present in more than one study, it was randomly assigned to one of them. This step was essential to avoid inflated results in the validation studies.

### Balanced Negative Sampling

A variant is considered to be statistically associated with a disorder if the *p*-value is smaller than 5 *×* 10^−8^. However, in GWAS, a class imbalance is typically present, where negative labels (i.e., no evidence of association) are more frequent. We address this by randomly selecting non-associated variants (*p*-values greater than 0.5), ensuring a balanced distribution of associated and non-associated variants in each chromosome.

### Annotation

Once the variants for model training are selected, we proceed with their annotation using the CADD file titled *“All possible Single nucleotide variations (SNVs) of GRCh38/hg38 incl. all annotations”*, available at https://cadd.gs.washington.edu/download, which contains precomputed scores and comprehensive annotations for all SNPs in the GRCh38/hg38 reference genome. CADD scores are excluded from the feature sets used to train the models.

### One Hot Encoding, Scaling and Duplicates Merging

All categorical variables are transformed using one-hot encoding, while continuous variables are standardised. To construct a unified data set comprising variants from all studies, we also apply one-hot encoding to the target variable, generating a binary column for each disease to indicate whether a variant is associated with that condition. Variants reported in multiple studies are consolidated into a single entry; therefore, variants significantly associated with more than one disorder retain positive labels for each relevant disease.

Variants with conflicting annotations for the same disease across studies were considered positives if they were associated in at least one study.

The CADD annotation pipeline can generate duplicate entries in the *Consequence* feature (i.e. the consequence of the variant on the transcript overlapping the genomic position), resulting in otherwise identical records that differ only in this categorical attribute. To resolve this, we one-hot encoded the *Consequence* feature and merged duplicate entries using the same procedure applied to the target variables.

### Missing Data

The CADD features mapping generated some missing values due to non availability of some features for certain genomic positions. All features with a missing rate higher than 40% were dropped; the remaining features are imputed with mean values for continuous features and ‘undefined’ for categorical features.

### 3 Demonstration, Modeling and Evaluation on Coronary Artery Disease

To demonstrate the practical utility of the proposed pipeline, we apply it to the prediction of genetic variants associated with CAD. Using the dataset generated through the pipeline, we design and evaluate a predictive model using a Leave-One-Group-Out Cross-Validation (LOGOCV).

To evaluate whether CADD-derived features can discriminate between CAD-associated and non-associated variants, we selected variants from GWAS conducted in individuals of European ancestry, with full summary statistics available for CAD (EFO 0001645; see Table 1). After running the described pipeline, we trained an XGBoost model and a Logistic Regression as baseline on the resulting dataset. To assess its performance, we implemented a LOGO-CV. In this settings the model were fine-tuned using Bayesian hyperparameter optimisation based on the Tree-structured Parzen Estimator, implemented via the Hyperopt library [9]. We defined the cross-validation folds based on GWAS study identified by their GCST ID. At each iteration, we held out all variants from two studies: one as a validation set for hyperparameter tuning and the other as an independent test set. This design ensures that the test data is never used during training or parameter optimisation, providing an unbiased estimate of model generalisation performance.

**Table 1.** CAD GWAS, showing total variants and class balance between significant and non-significant associations.

| GCST ID | Total SNPs | # associated SNPs | # non associated SNPs |
| --- | --- | --- | --- |
| GCST90132315 | 56,184 | 27,557 | 28,627 |
| GCST90132314 | 40,214 | 19,854 | 20,360 |
| GCST003116 | 4,623 | 2,203 | 2,420 |
| GCST000998 | 4,523 | 2,135 | 2,388 |
| GCST90309350 | 3,515 | 1,598 | 1,917 |
| GCST90274723 | 1,350 | 662 | 688 |
| **Total** | **110,409** | **54,009** | **56,400** |

Model performance was evaluated using the area under the receiver operating characteristic curve (AUROC) on the held-out test study. If a disease category includes *n* studies, this procedure yields *n* trained models, each associated with its own AUROC score and optimised hyperparameter set.

## 4 Results

The annotation produced a data set comprising 110,409 variants annotated with 187 features. The associations were divided into 54,009 significant and 56,400 non-significant associations.

Overall, significant and non-significant associations were balanced across chromosomes (Figure 1**B,C**), suggesting no strong chromosomal bias in the data set.

The analysis of feature correlations (Figure 1**D**) indicates that global collinearity is not prominent. The hierarchical correlation matrix reveals a clear block structure, where features cluster according to shared biological or technical functions. Clusters of highly correlated features as red blocks along the diagonal—highlight redundancy among annotations capturing similar biological signals. This pattern is particularly evident in the clustering of histone modifications (H3K27ac, H3K4me1, H3K4me3), which co-localise at active regulatory elements, and in conservation metrics (phyloP, phastCons), which jointly reflect evolutionary constraint. The feature correlation blocks identified indicate biologically coherent relationships shaped by shared genomic context and evolutionary constraints.

In the CAD variant classification task, the fine-tuned XGBoost model achieved an AUROC of 0.96 ± 0.04 (Figure 1**F**), substantially outperforming the baseline linear model. This improvement likely reflects the ability of XGBoost to capture the complex, non-linear relationships between genomic features and variant pathogenicity, as also suggested by the SHAP analyses. We further compared the model against the original CADD scores, which achieved an AUROC of 0.493, indicating near-random discrimination (Figure 1**G**). This marked performance difference is expected, as CADD was developed for a general variant deleteriousness prediction task rather than disease-specific CAD variant classification.

SHAP analysis (Figure 1**E**) further shows that the most influential features driving predictions are linked to frequency of mutation, particularly the density of SNPs in regions surrounding the variant, consistent with previous observations in the literature [10]. Notably, interactions with conservation-related metrics such as bStatistic (where lower scores reflect more conserved regions) and cHmm E4 (where higher scores indicate inactive chromatin) suggest that these mutational hotspots are preferentially associated with active, evolutionarily constrained genomic regions.

## 5 Conclusion

In this work, we developed a pipeline to generate an annotated database of variants associated with complex disorders. Its design allows users to retrieve a custom set of studies from the GWAS Catalog for any disorder available in the repository. The selected studies are automatically lifted to the latest genomic assembly, risk-associated variants are identified, and non-associated variants are generated through chromosome-stratified sampling, with their number matched to that of significantly associated variants within each chromosome. Variants are then extensively annotated with CADD-derived features, preprocessed, and one-hot encoded, yielding a database that is readily usable for downstream machine learning applications. We believe this framework opens the way for a wide range of bioinformatics and data science applications aimed at improving our understanding of the genetic architecture of complex diseases, allowing interpretable analyses of genomic risk predisposition, and ultimately supporting improved patient stratification. The proposed framework is not free from limitations. In particular, linkage disequilibrium (LD) was not explicitly modeled or controlled for in the current evaluation setting. Consequently, the reported predictive performance may also reflect the model’s ability to recognize loci associated with the trait. When variants in the training and test sets are correlated through shared LD structure or belong to the same associated loci, part of the observed predictive accuracy may arise from this shared genetic signal rather than from the model’s ability to generalize to independent genomic regions. Positives were defined as variants reaching genome-wide significance 5 *×* 10^−8^ with a positive effect estimate, and negatives as variants showing no evidence of association. Negatives were required to be clearly null rather than merely non-significant. Restricting the set to variants with no detectable signal and discarding the intermediate range of weakly associated variants, prevents underpowered true associations from being mislabeled as negatives. We further note that, for complex traits, purifying selection is weak to leave the constraint signal that clearly distinguishes pathogenic from benign variants. A set of true negatives therefore cannot be defined on evolutionary grounds, and the non-associated variants set we use represents the best available approximation. Future work will focus on several directions, including addressing current limitations such as LD correlations between variants, introducing an external validation on patient-specific data and expanding the feature set—particularly through the integration of scores derived from genomic foundational models such as Evo 2 [5] and evaluating the constructed datasets in downstream applications, ranging from comorbidity analysis to polygenic risk score modeling.

## 6 Conflict of interests

The authors declare no conflict of interest.

## Availability of data and software code

GWAS studies full summary statistics are available at https://ftp.ebi.ac.uk/pub/databases/gwas/summary_statistics/. CADD-annotated variants are available at https://cadd.gs.washington.edu/download. The code implementing the pipeline is available at https://github.com/freh-g/GWAnnotation.

## Notes

### Competing Interest Statement

The authors have declared no competing interest.

### Funding

- European Union Horizon 2020: 101136962
- UK Research and Innovation: 10098097, 10098097
- Swiss State Secretariat for Education, Research and Innovation (SERI)

